# Phosphatidylinositol phosphate binding domains exhibit complex dissociation properties at the inner leaflet of plasma membrane sheets

**DOI:** 10.1101/2021.02.17.431012

**Authors:** Madeline R. Sponholtz, Eric N. Senning

## Abstract

The pleckstrin homology (PH) domain is a lipid targeting motif that binds with high specificity to phosphatidylinositol phosphate (PIP) lipids. Using TIRF microscopy, we followed the dissociation of GFP-tagged PH domains from the plasma membranes of rapidly unroofed cells and found that AKT-PH and PLCδ1-PH dissociation kinetics can be distinguished by their effective k_off_ values determined from fitting fluorescence traces to a single exponential equation. Our measurements for the k_off_ of AKT-PH-GFP and PLCδ1-PH-GFP were significantly different (p < 0.05) at 0.39 ± 0.05 s^−1^ and 0.56 ± 0.17 s^−1^, respectively. Furthermore, we identified substantial rebinding events in our measurements of PLCδ1-PH-GFP dissociation kinetics. By applying inositol triphosphate (IP_3_) to samples during the unroofing process, we measured a much faster k_off_ of 1.54 ± 0.42 s^−1^ for PLCδ1-PH-GFP, indicating that rebinding events are significantly depressed through competitive action by IP_3_ for the same PH domain binding site as phosphatidylinositol (4,5)-bisphosphate (PIP_2_). We discuss the complex character of our PLCδ1-PH-GFP fluorescence decays in the context of membrane receptor and ligand theory to address the question of how free PIP_2_ levels modulate the interaction between membrane associated proteins and the plasma membrane.

## Introduction

Many essential cell signaling events occur at membrane surfaces, often involving the recruitment of intracellular signaling proteins to the membrane by conserved membrane targeting domains (1). The pleckstrin homology (PH) domain is the most common membrane targeting motif, and it serves as a critical element in numerous cell signaling pathways (2, 3). Although the majority of PH domains are general membrane scaffold modules, a subclass of PH domains bind with high specificity to phosphatidylinositol phosphate (PIP) lipids (3). For instance, the N-terminal PH domain of phosphoinositide phospholipase C-δ1 (PLCδ1) binds to phosphatidylinositol (4,5)-bisphosphate (PIP_2_) with a dissociation constant (K_d_) in the 1 to 10 μM range (2, 4–6). This binding in turn promotes activation of PLCδ1-mediated PIP_2_ hydrolysis (4). The PH domain of protein kinase B (AKT) can bind to either phosphatidylinositol (3,4)-bisphosphate (PI(3,4)P_2_) or phosphatidylinositol (3,4,5)-trisphosphate (PIP_3_), both of which are nominally present at a mere fraction of the PIP_2_ concentration in the plasma membrane (PM) (7, 8). In both AKT and PLC δ1, the PIP-specific PH domain serves to tether the enzymes’ catalytic activity to the PM.

Cell signaling is highly dynamic, and several methods have been utilized to measure the kinetics of PH domain interactions with lipids (2). Surface plasmon resonance (SPR) assay was used to measure a K_d_ of 2.8 μM for PLCδ1-PH-GFP binding to PIP_2_ (2.1 μM without GFP tagging) (6) and a K_d_ of 590 nM for AKT-PH binding to PIP_3_ (9). Dissociation rates (k_off_) have also been obtained from total internal reflection fluorescence (TIRF) measurements of a PH domain isolated from the general receptor for phosphoinositides, isoform 1 (GRP1) on a synthetic bilayer (10). In their study, the k_off_ of GRP1, which targets PIP_3_, ranged from 0.4 s^−1^ to 0.8 s^−1^ depending on the lipid environment (10).

Since SPR and fluorescence microscopy experiments with various PH domains have relied on artificial monolayer substrates or synthetic bilayers, we sought to assess the importance of PM architecture and composition to the k_off_ of PH domains by preserving the native lipid composition of the PM using PM sheets. There is broad appeal to using PM sheets as a biological experimental platform. Electron microscopists were the first to combine the preparative technique of obtaining PM sheets by unroofing with freeze-etch replica imaging to show detailed structures at the interface of the cytoplasm and the inner surface of the PM (11–13). The PM sheet permits structural studies of macromolecules residing on the inner leaflet of the PM because the cytoplasmic contents of the cell have been cleared from the preparation (14, 15). Moreover, access to intracellular binding sites of integral membrane proteins makes the PM sheet a useful preparation for optical microscopy-based structural studies of ion channels (16). Here we utilize ultrasonic unroofing, a process that disrupts all organelles and leaves a single PM sheet and its associated proteins intact. Once the cell is unroofed, the intracellular pool of fluorescently labeled molecules is rapidly shed, ensuring our TIRF signal is solely from membrane-associated proteins. Because evanescent illumination from TIRF drops off steeply only 200 nm from the coverslip, unroofing further limits the fluorescent contribution from the unbound pool of fluorescently labeled protein (17).

Here we implement cell unroofing to isolate a native PM bilayer and use TIRF microscopy to follow the subsequent dissociation of GFP-tagged PH domains from the membrane. Furthermore, we show that AKT-PH-GFP and PLCδ1-PH-GFP populations can be distinguished qualitatively when assessing fluorescence traces and quantitatively based on k_off_ obtained from fitting these data to single exponential equations. Additionally, in our PLCδ1-PH-GFP overexpression system, we observed rebinding events both in bulk and at the single molecule level. In applying inositol triphosphate (IP_3_) to samples during unroofing, we measured a more rapid k_off_ for PLCδ1-PH-GFP, suggesting that rebinding events in this experimental paradigm occur frequently enough to significantly depress k_off_ measurements. Our experimental findings reveal that rebinding may play an important role in PH domain interactions with the PM.

## Results

Although previous studies have assessed the dynamic interactions between phosphoinositides and PH domains, these have typically relied on artificial substrates or synthetic bilayers (2, 4–6). Because the composition of the membrane has been shown to influence the kinetics of such interactions, physiological insight could be derived from kinetic experiments conducted with PH domains and native PM (10). Here, we utilize cell unroofing to prepare native PM sheets and assess the subsequent dissociation of PH domains from the membrane.

### Plasma membrane sheets prepared by unroofing cells with ultrasound

The cell unroofing process begins with a series of solution changes to make adherent HEK293T cells susceptible to an ultrasonic pulse delivered by micro-tip ultrasonic probe. Our unroofing chambers are optimized for inverted epi-microscopy so that cells are visualized from underneath as a micro-tip sonicator probe is lowered into an open chamber and activated for the sonication step (Figure 1A). Following sonication, a region absent of cells is observed proximal to the location of the sonication probe. Regions of intact cells become prominent once more when the objective is positioned further away from the site of sonication (Figure 1E-G). To test whether the absence of intact cells closer to the sonicator probe was indicative of PM sheet preparation, we used the lipophilic dye DiI (ThermoFisher, Waltham MA) to stain the lipid bilayer of the PM. Membrane staining with DiI revealed a region of PM sheets that extended from directly underneath the sonicator probe to further removed locations where intact cells appear in differential interference contrast (DIC) images (Figure 1F,G).

**Figure 1.**
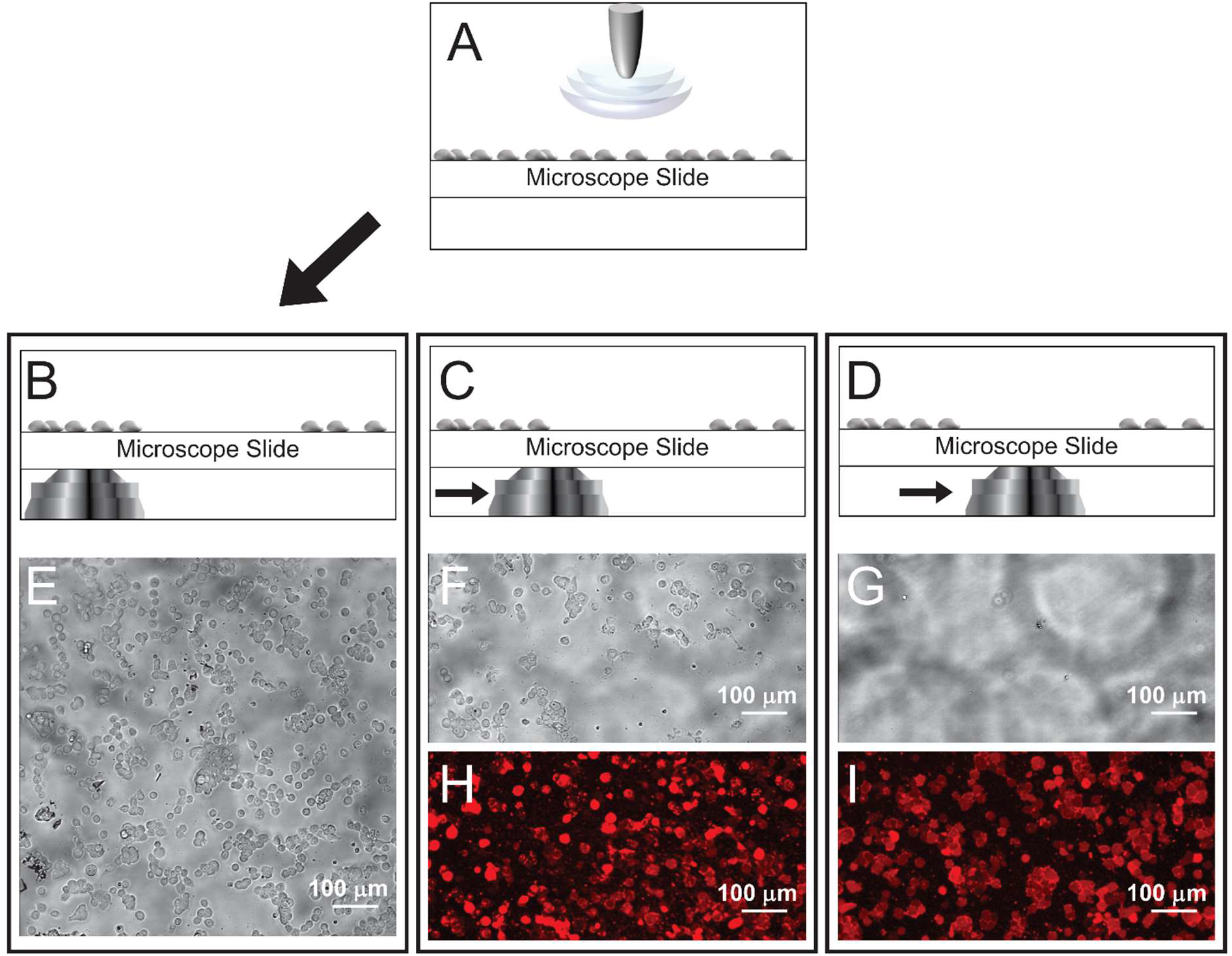
Unroofing HEK293T cells to form plasma membrane sheets. A brief sonication pulse “unroofs’’ the cells in a localized area (A). Cells are intact outside of the pulse range (B) and visible in a differential interference contrast (DIC) image (E), but within the pulse range (C,D), unroofing cells makes them disappear in DIC images (F,G). Lipophilic dyes, including Rhodamine B - C18 (R18) and DiI, can bind to the PM sheets that remain after unroofing and render them visible by fluorescence (H,I).

On closer inspection of the DiI stained PM sheets prepared by our unroofing method, we noted uneven fluorescence intensity on the peripheral edges of sheets formed from individual cells and at sites where neighboring adherent cells were in contact (Figure 1I). Heightened fluorescence intensities on the edges relative to the center of PM sheets are also observed with rhodamine B - C18 (R18, ThermoFisher, Waltham MA) labeling of the bilayer (Figure 2A). We then performed atomic force microscopy (AFM) scanning of unroofed PM sheet samples to assess whether the increased fluorescence intensity at the edges of DiI and R18 labeled sheets is due to incompletely unroofed regions. Using AFM, we found that the edges of PM sheets exhibited patches with a height that was approximately double that of the middle of a sheet (Figure 2B,C). Consistent with our hypothesis that the uneven fluorescence intensity is due to incompletely unroofed regions, the AFM results indicate that two bilayers remain along some of the edges in PM sheets.

**Figure 2.**
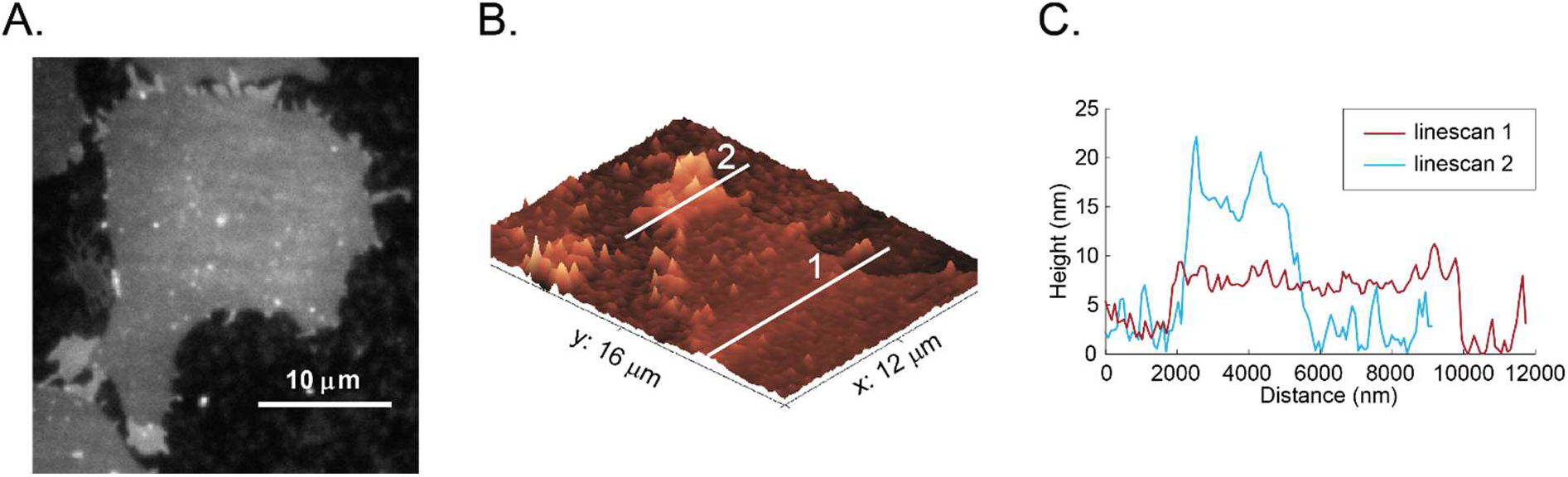
Characterizing unroofed HEK293T cells by fluorescence microscopy and AFM. (A) PM sheet loaded with R18 dye. Brighter regions on the periphery indicate partial unroofing where two bilayers are still present. (B,C) Atomic force microscopy (AFM) image of an unroofed cell on a glass coverslip confirms that most of the cell footprint is a single PM bilayer (mean thickness of higher region in linescan 1: 5.2 ± 2.1 nm), while thicker regions at the edges of cell footprints correspond to two bilayers (mean thickness of higher region in linescan 2: 12.9 ± 3.4 nm).

PM sheets prepared by unroofing exhibit a uniform thickness across large regions as demonstrated by lipophilic dye fluorescence and AFM measurements of the membrane height (Figure 2). Nevertheless, there is a concern that intracellular membranes remain attached to the PM sheet after unroofing, potentially contaminating our preparation. We estimated the degree to which intracellular membranes contaminate the PM sheets in our preparations by expressing the endoplasmic reticulum (ER) membrane marker Sec61b-GFP in HEK293T cells and monitoring the change in fluorescence upon unroofing. TIRF images of Sec61b-GFP fluorescence measured in intact cells and on PM sheets following unroofing show a sharp loss in this ER membrane marker (Supplemental Figure 1A). We confirmed the loss of ER membrane by examining unroofed HEK293T cells that express low levels of the fluorescently labeled ion channel transient receptor potential vanilloid type 1 (TRPV1-GFP). Prior to unroofing, the intact cells exhibited single molecule puncta of TRPV1-GFP near the edges of the cell footprint and a higher density of fluorescent molecules in the center of the cell (Supplemental Figure 1B). Following unroofing, the PM sheet shows a more even distribution of randomly positioned single molecule TRPV1-GFP (Supplemental Figure 1B). We interpret the decreased fluorescence in the center of the footprint after unroofing as the loss of ER membranes containing intracellular TRPV1-GFP. Numerous studies rely on TIRF microscopy to enhance the signal-to-noise ratio of fluorescent objects in the PM. Our pre- and post-unroofing experiments of Sec61b-GFP and TRPV1-GFP indicate that fluorescence from internal membranes contaminates signals from on or near the PM, despite the use of TIRF microscopy (Supplemental Figure 1). The marked absence of contaminating fluorescence signal in these samples following unroofing confirms that internal membranes are shed in the unroofing process and supports our contention that unroofing by sonication is a satisfactory method for preparing highly purified PM sheets.

Our method of sonicating cells to acquire PM sheets brought to light a concern that the biological activity of the PM could be lost during the unroofing step. Sonication is commonly used to disrupt cells during protein expression and purification methods for downstream biochemical assays, so we felt confident that the PM sheet sample retains most if not all of its enzymatic activity. We confirmed this by preparing PM sheets from cells co-transfected with TRPV1-SNAP and TRPV1-GFP and click-chemistry labeling the TRPV1-SNAP. In labeling the TRPV1-SNAP construct, the benzylguanine (BG) derivative of Dyomics dye DY-549P1 (DyL-549, Promega, Madison WI) is covalently attached to TRPV1-SNAP via an alkyltransferase reaction. Because TRPV1 forms an obligate tetramer, we expect partial colocalization between TRPV1-SNAP and TRPV1-GFP when both are expressed in the same cell. Activity of the SNAP enzymatic process revealed a consistent optical colocalization of TRPV1-SNAP labeled by DyL-549 and TRPV1-GFP in single molecule puncta (Supplemental Figure 2A). Overlap of fluorescent puncta was scored by optical colocalization (OC) measurements made with the JACoP plug-in of ImageJ (18), and OC scores were compared to randomized puncta distributed at the same density over the region of the PM sheet (Supplemental Figure 2A, right side). OC scores were considerably higher for TRPV1-SNAP and TRPV1-GFP compared to randomized puncta, confirming partial colocalization of the two TRPV1 constructs. As a point of comparison, we also determined the OC score of two membrane proteins that do not interact. In PM sheets derived from cells expressing tropomyosin receptor kinase A (TrkA)-SNAP and TRPV1-GFP, the OC score of red and green puncta was not markedly higher than a randomized sample (Supplemental Figure 2B).

### Dissociation kinetics of AKT-PH-GFP and PLCδ1-PH-GFP from plasma membrane sheets

Our characterization of PM sheets as experimental platforms that retain biochemical functionality persuaded us to investigate the interactions of pleckstrin homology (PH) domains with their known lipid targets. The PH domains of phospholipase-C δ1 (PLCδ1-PH) and AKT have been engineered into fusion proteins with GFP to serve as common lipid markers that target the phosphatidyl inositol headgroups of PI(4,5)P_2_ and PI(3,4,5)P_2_, respectively. Consequently, the dynamic changes of these lipids in cellular membranes may be monitored by fluorescence microscopy (19, 20). Previous SPR experiments on purified PLCδ1-PH have measured its affinity to and dissociation from PI(4,5)P_2_ enriched supported lipid monolayers (6). We sought to compare the dissociation kinetics measured in these SPR experiments to the k_off_ of PLCδ1-PH-GFP from PM sheets prepared by cell unroofing.

To measure AKT-PH-GFP and PLCδ1-PH-GFP kinetics, we followed our standard procedure of using a series of swelling solutions to destabilize cell integrity prior to sonication. HEK293T cells heterologously expressing the GFP-tagged PH domain constructs were imaged by TIRF microscopy during and after unroofing. To confirm that the unroofing step generated a PM sheet, we followed each experiment with DiI labeling to fluorescently visualize the PM bilayer (Figure 3A). Each unroofing experiment captured the ensemble dissociation kinetics of either AKT-PH-GFP or PLCδ1-PH-GFP from the freshly exposed inner leaflet of the PM sheet, and the fluorescence loss in a region of interest (ROI) was fit to a single or double exponential equation (Figure 3B, Supplemental Movies 1,2). The fits of AKT-PH-GFP and PLCδ1-PH-GFP dissociation were slightly improved with increased parametrization from a single exponential to a double exponential (Figure 3B). Nevertheless, because the general feature of the fluorescence decay in all experiments could be adequately captured with a single exponential function, we elected to compare effective dissociation constants (k_off_) derived from single exponential fits of PLCδ1-PH-GFP and AKT-PH-GFP fluorescence decay data (Figure 3C, Table 1).

**Figure 3.**
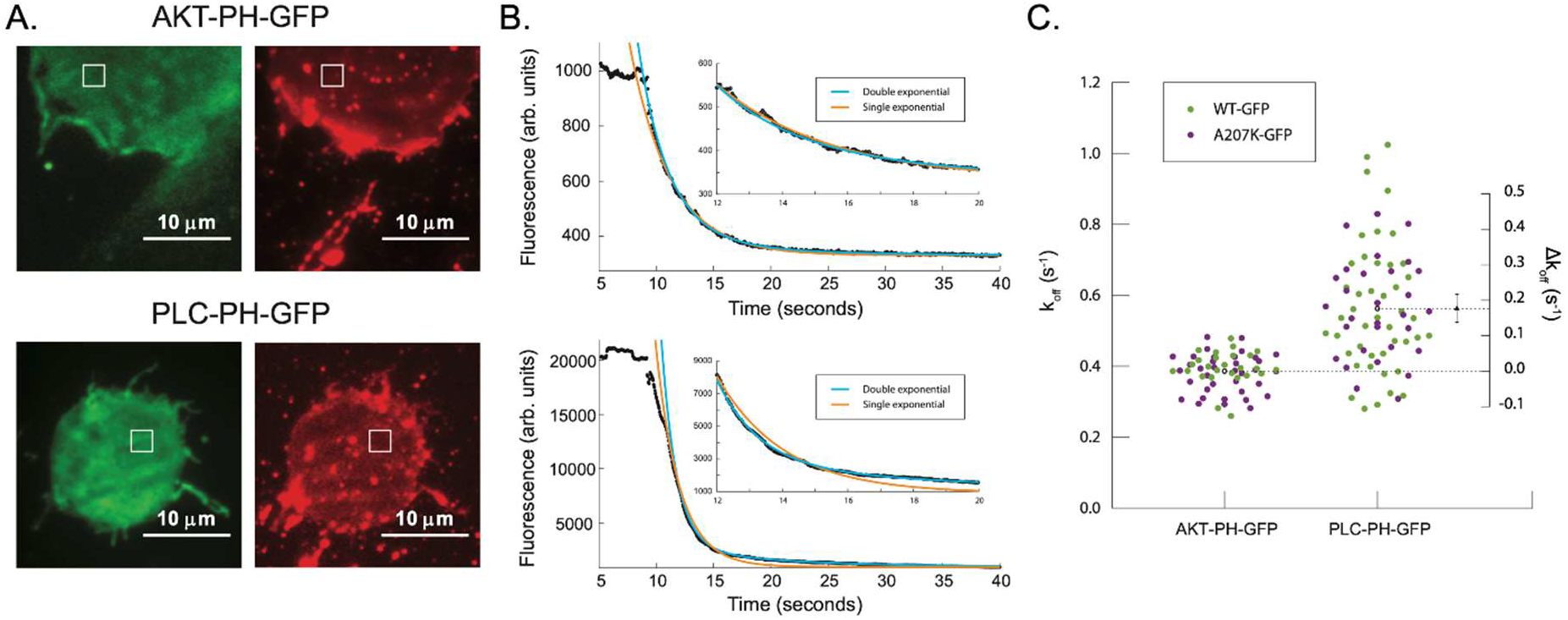
PLCδ1-PH-GFP and AKT-PH-GFP can be distinguished based on k_off_. (A) Representative images of HEK293T cells expressing either AKT-PH-GFP or PLCδ1-PH-GFP with white boxes demarcating selected regions of interest (ROI). Cells were excited at 488 nm in TIRF to capture PH-GFP dissociation (left, Supplemental Movies 1,2) and excited at 561 nm in TIRF to confirm the presence of PM sheets with DiI (right). (B) Representative fluorescence traces specific to the regions of interest (top, AKT-PH-GFP; bottom, PLCδ1-PH-GFP. Cells were unroofed approximately 10 s into each recording, with fluorescence drop-off corresponding to dissociation of PH domains from the membrane. Fluorescence traces were fitted to double (cyan) and single (orange) exponential functions. Inset displays details of fitting to fluorescence data. (C) Aggregate data of single exponential decay rates observed in cells expressing PH domains tagged with different GFP variants. WT AKT-PH-GFP k_off_ = 0.38 ± 0.05 s^−1^, AKT-PH-GFP(A207K) k_off_ = 0.40 ± 0.05 s^−1^, pooled AKT-PH-GFP k_off_ = 0.39 ± 0.05 s^−1^, WT PLCδ1-PH-GFP k_off_ = 0.56 ± 0.18 s^−1^, PLCδ1-PH-GFP(A207K) k_off_ = 0.56 ± 0.14 s^−1^, pooled PLCδ1-PH-GFP k_off_ = 0.56 ± 0.17 s^−1^. Ordinate axis on the right indicates difference of the mean k_off_ values and error bars represent a

**Table 1.**
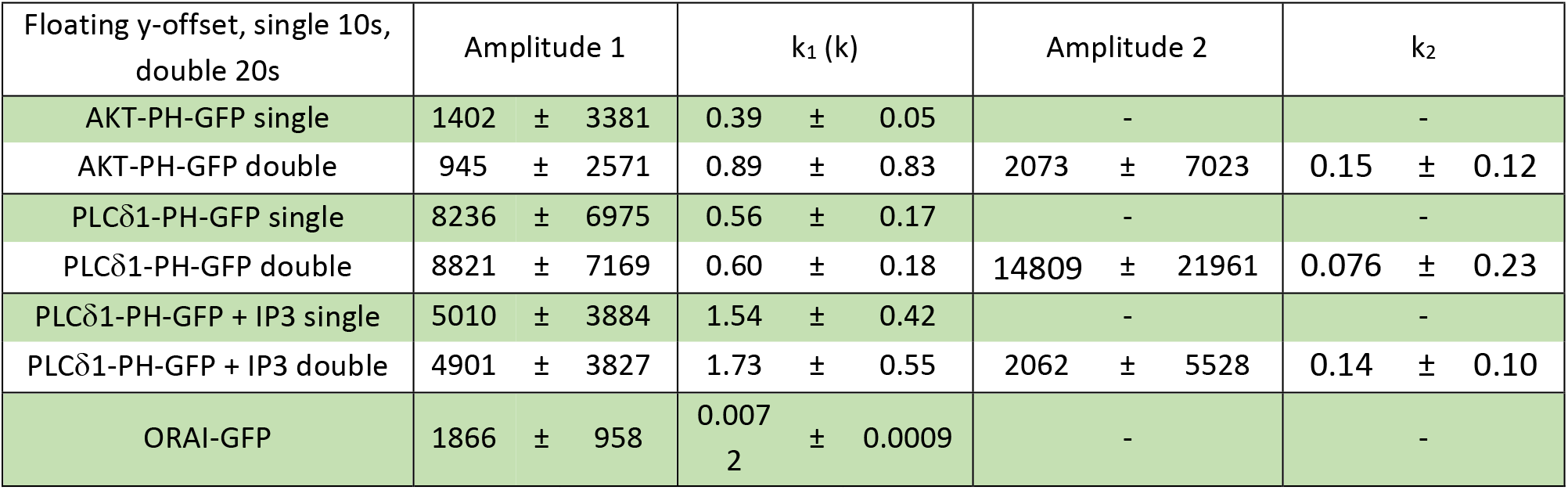

In assessing the source of the slow component of our fluorescence decay data (Table 1), we considered the possibility that the GFP portion of the PH domain constructs may induce the formation of a short-lived GFP-GFP dimer, which could affect the dissociation kinetics from the PM (21, 22). To address this concern, we repeated the dissociation experiments with constructs containing the dimerization deficient form of GFP with an A207K mutation (Zacharias et al., 2002). Switching to A207K-GFP did not appreciably alter the effective k_off_ of PLCδ1-PH-GFP(A207K) (Figure 3C). However, there was a small but significant change observed in the k_off_ of AKT-PH-GFP(A207K) in comparison to that of the wild-type construct, AKT-PH-GFP. This small difference between the k_off_ of AKT-PH-GFP and AKT-PH-GFP(A207K) did not detract from the effect size of dissociation kinetics measured from pooled (wild-type and A207K GFP) AKT-PH-GFP and PLCδ1-PH-GFP constructs, with PLCδ1-PH-GFP displaying a significantly faster k_off_ in comparison to that of AKT-PH-GFP (Figure 3C). For the remainder of this study, we will refer to pooled data from wild-type and A207K-GFP constructs.

To test the possibility that the slow component of the fluorescence decay stems from the photobleaching of GFP under our imaging conditions, we recorded fluorescence decays in unroofed HEK293T cells that heterologously expressed the integral membrane protein Orai1-E106A-GFP. As the GFP tag is PM bound in these unroofed cells, the fluorescence loss is expected to faithfully reflect the rate at which GFP is photobleached with our imaging system. The mean rate of fluorescence loss for Orai1-E106A-GFP in PM sheets was 0.0072 +/− 0.0009 s^−1^ (Supplemental Figure 3). Photobleaching of the GFP chromophore was assumed to occur independent of PH domain dissociation, so we expected a double exponential would describe dissociation and photobleaching rates for faster and slower constants, respectively. However, the Orai1-E106A-GFP photobleaching rate was not in close agreement with either of the slow components obtained from double exponential fits of PLCδ1-PH-GFP or AKT-PH-GFP (Table 1). Both AKT-PH-GFP and PLCδ1-PH-GFP constructs exhibited complicated fluorescence decay kinetics, and when fit with a double exponential function the slower decay constant could not be construed as fluorophore photobleaching or as a consequence of GFP dimerization.

### Unroofed HEK293T cells acquire PLCδ1-PH-GFP from neighboring transfected cells

In our attempts to determine the origin of the complicated decay kinetics in PLCδ1-PH-GFP dissociation from unroofed cells, we observed PLCδ1-PH-GFP rebinding events on PM sheets originally devoid of PLCδ1-PH-GFP. A representative example of this occurrence is shown in Figure 4, in which two cells are imaged side-by-side before (Figure 4A left and right panels, Supplemental Movie 3), during (Figure 4B), and after unroofing (Figure 4A center panel). The fluorescence reported for the demarcated ROI in the footprint of the untransfected cell is initially low (Figure 4B left panel and trace in Figure 4C before leftmost arrowhead). Upon unroofing, PLCδ1-PH-GFP that is lost from the neighboring, transfected cell begins to accumulate on the newly exposed PM sheet from the untransfected cell, as indicated by a rise in the fluorescence trace in Figure 4C (region between leftmost arrowhead and middle arrowhead). As the local concentration of PLCδ1-PH-GFP in the buffer decreases due to constant buffer exchange in the chamber, the rate of PLCδ1-PH-GFP acquisition in the untransfected membrane drops below the rate of PLCδ1-PH-GFP loss, resulting in a net loss of fluorescence (Figure 4B right panel and Figure 4C rightmost arrowhead). We interpreted the rise and fall of fluorescence on the untransfected PM sheet as PLCδ1-PH-GFP rebinding after its initial release from the neighboring, transfected cell.

**Figure 4.**
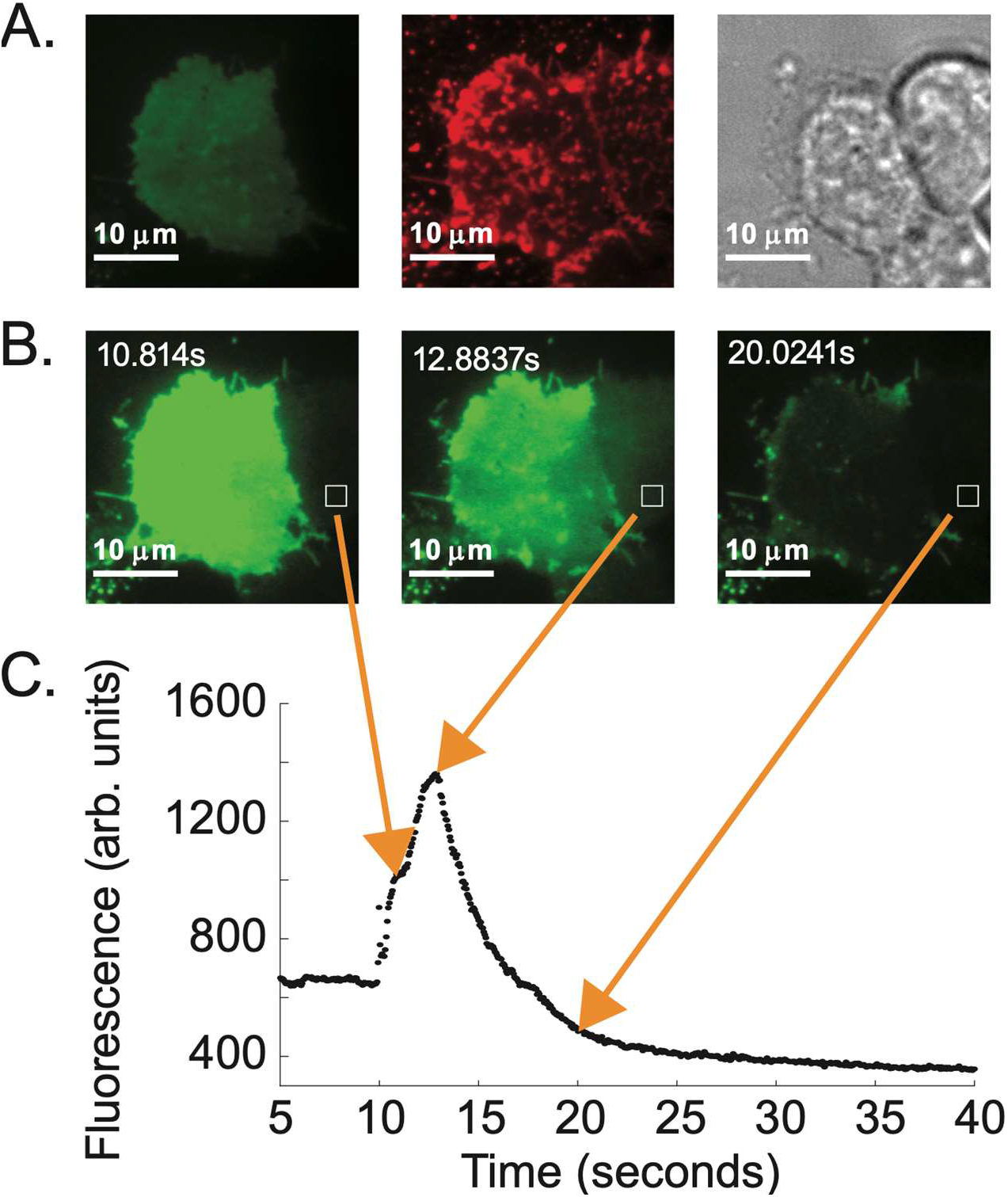
PLCδ1-PH-GFP rebinding occurs in unroofed HEK293T cells. (A) Representative images of cells transfected with PLCδ1-PH-GFP with white boxes demarcating selected ROIs. Cells excited at 488 nm prior to unroofing show one cell expressing PLCδ1-PH-GFP (left). DIC imaging shows two additional cells that failed to express the construct are present within the frame (right). Following unroofing, cells treated with DiI and excited at 561 nm confirm the presence of PM sheets from three cells (center). (B) After unroofing, free PLCδ1-PH-GFP diffuses from the transfected cell and binds to target lipids on nearby ‘recipient’ PM sheets from cells that failed to express the construct (see Supplemental Movie 3). (C) Representative trace specific to the selected ROI demarcated by the white boxes in (B).

Rebinding untransfected PM sheets occurs across a length-scale on the order of the cell width, and it stands to reason that PLCδ1-PH-GFP rebinding similarly happens within the same PM sheet that experiences a net loss of PLCδ1-PH-GFP following the unroofing event. To evaluate PLCδ1-PH-GFP rebinding at the single molecule level, we unroofed HEK293T cells expressing PLCδ1-PH-GFP and continuously monitored single molecule binding events of PLCδ1-PH-GFP to the PM sheet (23). The perfusion of fresh buffer to the chamber was switched off immediately after unroofing, and the liberated PLCδ1-PH-GFP from unroofed cells was permitted to equilibrate within the chamber volume so that infrequent binding events of PLCδ1-PH-GFP to the PM sheet could be observed. Optical recordings of single PLCδ1-PH-GFP molecules show that binding events occur at random intervals and indiscriminately across the PM sheet (Figure 5 and Supplemental Movie 4). Fluorescent features move randomly on the surface of the PM sheet as would be expected for PLCδ1-PH-GFP when interacting with lipids in a fluid membrane. When a final concentration of 100 μM IP_3_ is added to the bath at the 6 second mark in the movie, individual fluorescent events no longer occur (Supplemental Movie 4). We also note that fluorescent features which moved in the lead-up to IP_3_ addition were rapidly lost, and a smaller subset of static fluorescent sites remained for a longer duration on the sheet or photobleached in accordance with non-specific fluorescent interactions (Figure 5D-F). We interpret the loss of fluorescence events as IP_3_ binding to the PLCδ1-PH-GFP pocket for PIP_2_, rendering PLCδ1-PH-GFP in solution within the perfusion chamber unavailable to interact with the PM sheet.

**Figure 5.**
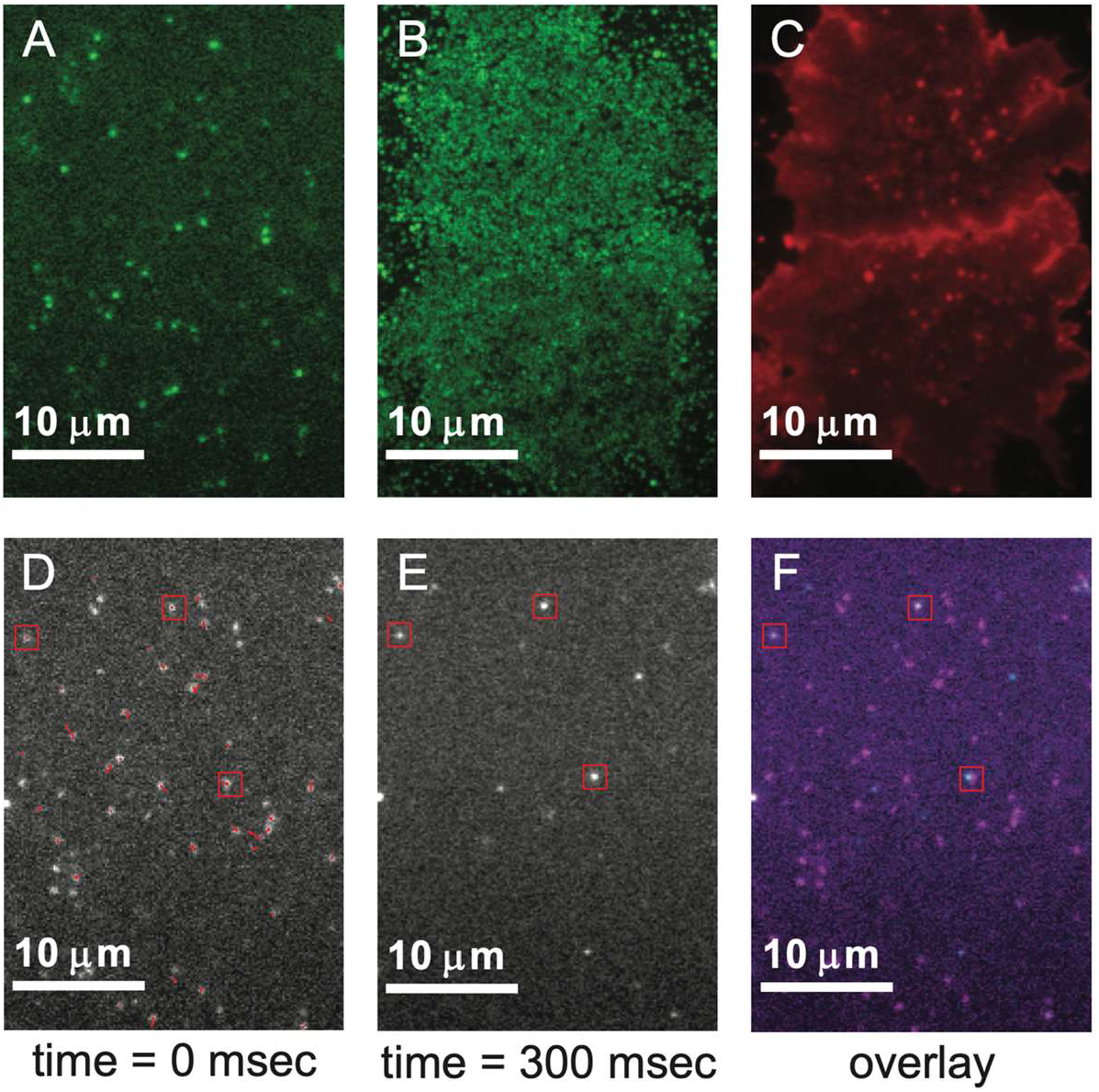
Addition of IP_3_ prevents PLCδ1-PH-GFP rebinding. (A) Single frame in movie of PLCδ1-PH-GFP molecules transiently binding unroofed HEK293T cells (see Supplemental Movie 4). (B) 800 frame movie projection demonstrates the range of movement for PLCδ1-PH-GFP as these fluorescently-tagged molecules traverse the PM sheets. (C) DiI image confirms the presence of PM sheets following unroofing. (D) Overlay of movie frame just prior to IP_3_ addition with tracks of moving PLCδ1-PH-GFP before unbinding. Red boxes demarcate non-specific signal from static positions. (E) After IP_3_ addition (for a total concentration of 100 μM IP_3_), the majority of PLCδ1-PH-GFP signal is lost, with the exception of non-specific signal (red boxes). (F) Overlay of movie frames shown in D (magenta) and E (cyan) with static positions indicated (red boxes).

The acute sequestration of PLCδ1-PH-GFP from PM sheets through the addition of IP_3_ observed at the single molecule level prompted us to incorporate IP_3_ in our unroofing experiments to measure PLCδ1-PH-GFP dissociation kinetics. With the addition of 100 μM IP_3_ to the perfusion buffer during the sonication step (see methods for buffer details), we once again measured the k_off_ of PLCδ1-PH-GFP from HEK293T cells (Figure 6, Supplemental Movie 5). Under identical unroofing conditions as described for the experiments in Figure 3, we measured a k_off_ for PLCδ1-PH-GFP in 100 μM IP_3_ buffer (PLCδ1-PH-GFP/IP_3_) that was more than twice the rate of that without IP_3_ (Figure 6C). As was the case for PLCδ1-PH-GFP alone, visual assessment of fitting the PLCδ1-PH-GFP/IP_3_ experimental data to a double exponential improved in comparison to a single exponential function (Figure 6B). Nevertheless, a distinguishing feature of the decays emerges in the PLCδ1-PH-GFP/IP_3_ fluorescence decay data which is not apparent in the PLCδ1-PH-GFP data. A semi-log plot of the PLCδ1-PH-GFP/IP_3_ fluorescence decays reveals a transition occurring at around 1 s, which demarcates the end of a period of faster dynamics and the beginning of slower dynamics in the fluorescence measurements (Figure 6D). The absence of such a clear transition between two dynamic regimes in the PLCδ1-PH-GFP unroofing experiments without IP_3_ and the significant overall increase in effective k_off_ for PLCδ1-PH-GFP in the presence of 100 μM IP_3_ lead us to believe that PLCδ1-PH-GFP rebinding events are a frequent occurrence in the context of our unroofing experiments. Moreover, the persistence of a slow dynamic process in PLCδ1-PH-GFP/IP_3_ experiments suggests that loss of PLCδ1-PH-GFP to near infinite dilution cannot effect a simple rate of PLCδ1-PH-GFP depletion from the native PM at our experimental timescale.

**Figure 6.**
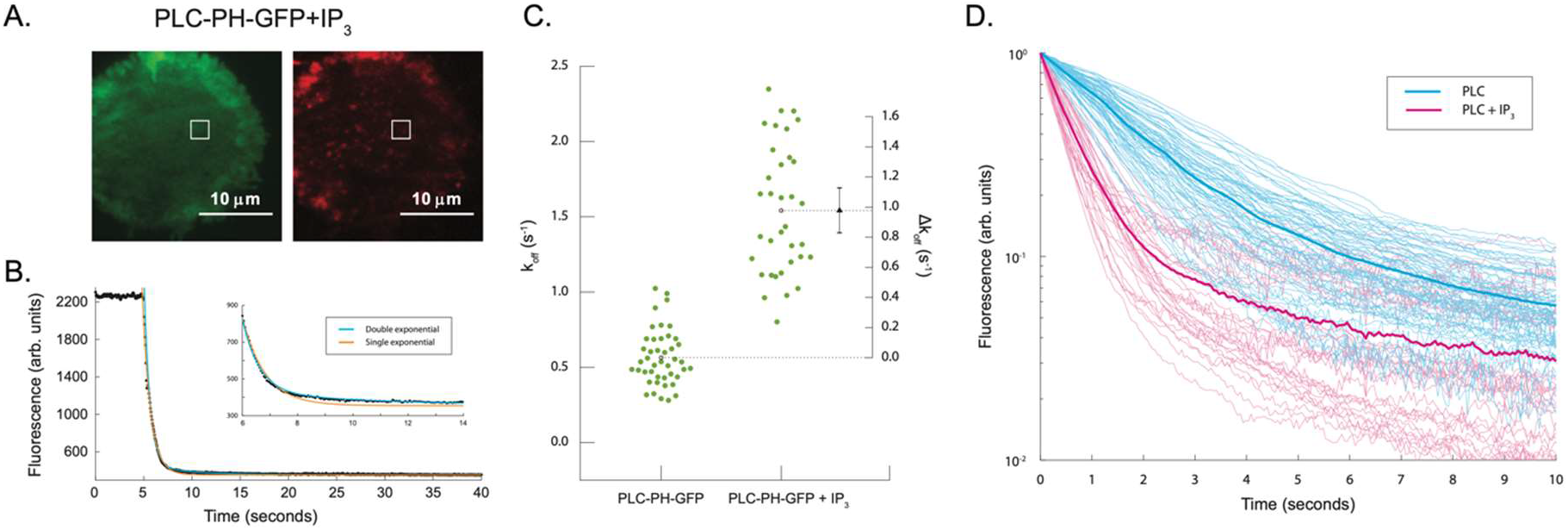
Measured rates of PLCδ1-PH-GFP dissociation increase in the presence of IP_3_. (A) HEK293T cells expressing PLCδ1-PH-GFP were excited at 488 nm to follow PLCδ1-PH-GFP dissociation (left, Supplemental Movie 5) and excited at 561 nm following DiI treatment to confirm the presence of PM sheets (right). White boxes demarcate the selected ROI. (B) Representative fluorescence trace specific to the ROI. Cells were unroofed around 5 s into each recording, with fluorescence drop-off corresponding to dissociation of PLCδ1-PH-GFP from the membrane in the presence of 100 μM IP_3_. Fluorescence traces were fitted to double (cyan) and single (orange) exponential functions. Inset displays details of fitting to fluorescence data. (C) Aggregate data from cells expressing PLCδ1-PH-GFP treated with vehicle or 100 μM IP_3_ fitted to a single exponential function. PLCδ1-PH-GFP k_off_ = 0.56 ± 0.17 s^−1^ and PLCδ1-PH-GFP with IP_3_ k_off_ = 1.54 ± 0.42 s^−1^. (D) All fluorescence traces for PLCδ1-PH-GFP treated with vehicle or IP_3_ normalized to their maximum value following unroofing and displayed on a semi-log y-axis plot. Thick lines represent averages of individually normalized data from the respective PLCδ1-PH-GFP (blue) and PLCδ1-PH-GFP + IP_3_ (magenta) sets.

## Discussion

As recently shown, we unroof cells on an inverted microscope setup by immersing a Branson sonifier microtip in an optically accessible perfusion chamber (24). Our sonication technique for unroofing disrupts the HEK293T cell, clearing away the organelles and cytosolic material by continual perfusion and retaining a single bilayer of PM, which we refer to as the PM sheet (Figure 2 and Supplemental Figure 1). We also confirmed the biological activity of the integral membrane proteins retained in the PM sheet by labeling single molecules of TRPV1-SNAP with the click-chemistry dye DyL-549 and performing optical colocalization with TRPV1-GFP (Supplemental Figure 2). The rapid sonication step and accessibility of the sample by fluorescence microscopy motivated us to investigate kinetics of the PH domains from PLCδ1 and AKT by cell unroofing.

The PH domains of PLCδ1 and AKT were fused to GFP and expressed in HEK293T cells to measure their respective dissociation kinetics from the inner leaflet of the PM by TIRF microscopy during cell unroofing. We measured an effective k_off_ for PLCδ1-PH-GFP that was around 50% greater than that obtained for AKT-PH-GFP. Our experimental data for both constructs were more optimally fit by a double exponential, but we felt that a single exponential function could adequately capture the significant changes to the dynamics observed between AKT-PH-GFP and PLCδ1-PH-GFP. Several control experiments were conducted to rule out the possibility that a confounding component of the decay was either an artifact due to GFP dimerization (Figure 3C) or a result of GFP photodegradation during the experiment (Supplemental Figure 4), and it could not be attributed to either of these. Nevertheless, a small effect of the dimerization deficient A207K mutant GFP compared to wild-type GFP was observed in the AKT-PH-GFP data, demonstrating the sensitivity of this technique to small perturbations in our experiment (Figure 3C).

Further inquiry into the nonmonotonic decay of PLCδ1-PH-GFP and AKT-PH-GFP fluorescence revealed that dissociating molecules of PLCδ1-PH-GFP originating in a transfected cell are capable of rebinding to a neighboring unroofed cell that was not transfected with the construct (Figure 4). PH domain rebinding has been previously observed with TIRF microscopy in studies on a representative PH domain isolated from GRP1 with a synthetic bilayer (10). Since rebinding is a phenomenon that can reach across the dimensions of a cell during our experiment, we conclude that rebinding also occurs within the footprint of a single cell. Thus, the time-dependent, net loss of PLCδ1-PH-GFP from an unroofed cell is a summation of terms that represent: 1. dissociation kinetics of the bound component and 2. rebinding kinetics of a recently unbound component.

Our unroofing experiments with IP_3_ demonstrated that our measured effective k_off_ is highly dependent on the availability of the PIP_2_ binding pocket in PLCδ1-PH-GFP (Figure 6). We surmise from these data that PLCδ1-PH-GFP is first released from the PM in a stochastic manner consistent with first order rate kinetics; however, in the absence of saturating IP_3_, the PLCδ1-PH-GFP binding pocket is then available to rebind the PM. Although the complicated kinetics of PLCδ1-PH-GFP rebinding are reduced if IP_3_ competes for the PIP_2_ headgroup, a slow kinetic component remains in our IP_3_ unroofing data. We favor the interpretation that the remaining slow component persists under high concentrations of IP_3_ due to competition between IP_3_ and PIP_2_ for the same binding pocket.

Our method to measure the effective k_off_ for PH domains from unroofed PM sheets bears similarity to SPR kinetic measurements. Loss of the cytosolic pool of PLCδ1-PH-GFP or AKT-PH-GFP with cell unroofing is analogous to the rapid concentration jump in analyte that is used to measure ligand-analyte interactions by SPR. Additionally, the response units (RU) measured in SPR correspond to our detection of fluorescence in the PM sheet. One critical aspect that differs between these methodologies is the geometry and chemistry of our surface bound ligand. In a study by Hirose and co-authors, the PLCδ1-PH domain interacts with PIP_2_ in a supported monolayer comprised of phosphatidylcholine and 3% PIP_2_ (6). Our unroofing experiments replicate the geometrical constraints of PLCδ1-PH bound to its interaction partner PIP_2_ on the 2-D fluid inner leaflet of the PM. Furthermore, PH domain kinetics in unroofing experiments measure interactions with native partners in the native PM environment. Our kinetic measurements of PLC1-PH-GFP dissociating from the PM demonstrate that the additional presence of IP_3_ has a significant impact on the k_off_. This observation may be of practical importance when probing cellular phosphoinositide dynamics during GPCR stimulation. In the absence of IP_3_, there is a pronounced reluctance of the PLC1-PH-GFP domain to dissociate from the membrane which is relieved only if ample IP_3_ is washed onto the unroofed cell. We interpret these measurements to mean that a simplified 1-step kinetic scheme between PIP_2_ bound by PLCδ1-PH and cytosolic PLCδ1-PH in the intact cell should be reconsidered. Moreover, dynamic models of phosphoinositide levels in cellular membranes have relied on biophysical measurements by Hirose et al. and time resolved fluorescence signals from PLCδ1-PH as well as other lipid-specific probes, and we propose that further characterization of the probe kinetics with native PM sheets will sharpen the precision and predictive capacity of such models (25–27).

Given the differences between our cell unroofing experiments and dissociation measurements of PLCδ1-PH done with SPR in the presence of IP_3_, our respective k_off_ values of 1.54 s^−1^ and 5.25 s^−1^ (6) are, nevertheless, fairly close. However, by maintaining the topology and native membrane environment in our unroofing experiments, we were able to broach the question of whether rebinding is meaningful in the context of the cellular environment. Surface receptor and ligand binding events have been the object of intense study over a long period of time. Berg and Purcell provided early theoretical evidence that diffusive behavior of ligand near the cell surface could ensure repeated encounters between ligand and receptor even at low receptor densities, and a broad literature on the subject has grown over time, extending the principal ideas to greater complexity (28-32). In a simplified kinetic scheme for ligand (L) binding and unbinding to a receptor (R),

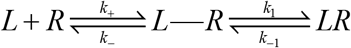

where (*k*_+_, *k*_−_) refer to the diffusional kinetic constants for an encounter complex (L-R) and (*k*_1_, *k*_−1_) relate to the chemical kinetic rates for LR, the overall association and dissociation rates can be formulated as 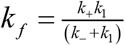 and, 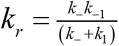, respectively. The overall dissociation rate (*k_r_*) would reduce to *k*_−1_ if *k*_−_ ≫ *k*_1_, and a monotonic exponential decay would be observed in kinetic experiments given this scenario. Neither PLCδ1-PH-GFP nor AKT-PH-GFP data from our unroofing experiments exhibited characteristic single exponential decays that would reflect *k*_−_ ≫ *k*_1_. Rather, we observed non-monotonic decays with both PH domains, and in the case of PLCδ1-PH-GFP, the addition of 100 μM IP_3_ significantly altered the overall dissociation rate. To explain these data, we turn to Delisi and Wiegel’s argument that a shift from *k*_−_ ≫ *k*_1_ to *k*_−_ ≈ *k*_1_ introduces a strong perturbation from the diffusion controlled dissociation rate (*k*_−_), and this gives rise to an increased complexity in the overall dissociation rate (*k_r_*). Recovery of simpler exponential decays by sequestration of PLCδ1-PH-GFP with IP_3_ provides further evidence that the diffusion-controlled dissociation rate (*k*_−_) plays an important role in the overall dissociation rate *k_r_*.

Previous determinations of the *K_D_* for PLCδ1-PH with membrane lipids by various methods noted complex behaviors ascribed to non-specific electrostatic and hydrophobic interactions (33, 34). We elected to model PLCδ1-PH-GFP dissociation kinetics from the PM as a process that separates the encounter complex from a diffusion controlled step (*k*_−_) because the latter term may incoporate energetic terms for non-specfic interactions with the PM. Delisi and Wiegel derive a convenient expression for the flux of the ligand departing the receptors on the cell membrane, *J*_−_, which may be equated to the diffusion limited off rate,

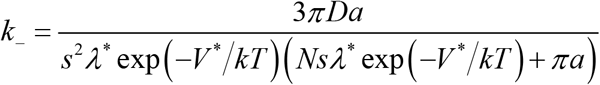

where D is the diffusion coefficient, N is the number of receptors on the cell membrane, s is the radius of a disc representing the footprint of the receptor, a is the radius of the cell represented as a sphere, and *λ*\* exp(−*V*\*/*kT*) represents an energy term for non-specific interactions. Non-specific interaction energies between the ligand and PM are thus accounted for in *J*_−_ and result in constant, concentration-independent terms for *k*_−_. In the absence of a strong perturbation through non-specific interactions, we deduce that the following expression holds for our system at large *a* relative to *s*when the surface of the cell may be approximated by a plane,

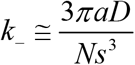

Thus, *k*_−_ varies inversely with the number of free receptor sites, *N*, which we interpret as free PIP_2_. In our unroofing experments, dissociation kinetics of PLCδ1-PH-GFP in the presence of 100 μM IP_3_ are separable into a fast regime immediately after unroofing and a slow regime at later times (Figure 6D). Keeping to an interpretation that PLCδ1-PH-GFP occupies a significant amount of the PIP_2_ during the early phase, we hypothesize that IP_3_ outcompetes PIP_2_ and quenches newly released PLCδ1-PH-GFP. After several seconds, during the slow regime, IP_3_ is still available to bind newly released PLCδ1-PH-GFP, but the free concentration of PIP_2_ is now elevated to compete with IP_3_ for the PLCδ1-PH-GFP binding pocket, which alters the circumstances from *k*_−_ ≫ *k*_1_ to *k*_−_ ≈ *k*_1_ and ensures a reduced overall dissociation rate, *k_r_*, at later times.

At this point it may be worth asking what the significance is in observing a dependence on *k*_−_ for PH domain dissociation kinetics from native PMs. First, unlike SPR experiments, which rely on reconstituted monolayers, our experiments are conducted on native PM sheets. This provides a greater understanding of how the unique 2-D architecture and lipid composition of the PM contribute to the interactions between PH domains and their target lipids. Second, the importance of the diffusion-controlled dissociation rate (*k*_−_) as a limiting step in the overall dissociation rate (*k_r_*) of PLCδ1-PH-GFP from PIP_2_ in the PM cannot be overlooked. Receptor and ligand rebinding events are tied to *k*_−_, which is inversely proportional to the number of available receptor sites. Since free PIP_2_ is known to be dynamically maintained as a second messenger in the PM, it stands to reason that *k*_−_ is an active homeostatic mechanism for any membrane-associated protein that specifically binds this phosphoinositide. Moreover, this property is likely shared with AKT and other phosphoinositide binding proteins, in which the potential to rebind increases concomitant with the concentration of free phosphoinositides.

## Methods

### Imaging and Cell Unroofing

We followed a previously reported unroofing protocol with minor modifications (24). HBR, stabilization buffer (SB, 70 mM KCl [Sigma-Aldrich, St. Louis MO], 30 mM HEPES [Sigma-Aldrich, St. Louis MO], and 1 mM MgCl_2_ [ThermoFisher, Waltham MA]; initial pH measured at 5.2 and adjusted to 7.4 with KOH [ThermoFisher, Waltham MA]; filter sterilized through a 0.45-μm membrane), swell buffer (a dilution of one part stabilization buffer to two parts water), and filter sterilized 0.1 mg/ml poly-L-lysine hydrobromide (P1274, MW 70,000-150,000, Sigma-Aldrich, St. Louis MO) in PBS were prepared prior to unroofing. After cells were passaged onto coverslips, they were incubated for 2–4 hours at 37°C in HBR and then placed in our perfusion chamber on the Eclipse Ti-E inverted total internal reflection fluorescence (TIRF) microscope (Nikon Instruments, Melville NY) at room temperature. Gentle flow was used to wash with HBR for 5-10 min. The sonicator probe (SFX250 Sonifier with 1/2” horn and 3 mm tapered microtip, Branson Ultrasonics, VWR International, Radnor PA) was then positioned in the chamber with its tip 2 mm above the coverslip. Poly-L-lysine was then perfused through the chamber for 10 s. Treatment with poly-L-lysine caused the cell footprint to expand and improved adherence to the coverslip upon sonication. Swell buffer was then perfused for 10 s, and then stabilization buffer was used to rinse the chamber. Imaging at 488 nm with the microscope commenced after rinsing for 20 s. After an additional 10 s of rinsing, the sonicator probe was used to give a 0.4 s sonic pulse with a power setting of 2. The cells were imaged for an additional 30 s, after which the probe was removed from the chamber and unroofing was confirmed through DIC imaging. Final confirmation of unroofing was by application of 1 nM 1,1’-Dioctadecyl-3,3,3’,3’-Tetramethylindocarbocyanine Perchlorate (DiI, Invitorgen, ThermoFisher, Waltham MA) in stabilization buffer, followed by imaging at 561 nm. DiI was made as a 1-mM stock in DMSO (Sigma-Aldrich, St. Louis MO) and stored at −20°C.

Dissociation rate measurements for PLCδ1-PH-GFP in 100 μM IP_3_ buffer were conducted as described earlier with the following adjustments. 100 μM IP_3_ in stabilization buffer was perfused through the chamber 10 s prior to unroofing. Imaging commenced 5 s prior to unroofing, and sonication proceeded at the normal time point in the sequence of buffer exchanges. Perfusion was switched back to stabilization buffer without IP_3_ 10 s after unroofing and the cells were imaged for an additional 20 s, after which the probe was removed from the chamber and unroofing was confirmed as described earlier.

Cell unroofing experiments were performed using an Eclipse Ti-E inverted TIRF microscope with a 100X oil immersion objective with the Perfect Focus system (Nikon Instruments, Melville NY) for preventing focal drift. All imaging data were collected with NIS-Elements AR software (Nikon Instruments, Melville NY) controlling an Andor iXon Life 897 camera (Oxford Instruments, UK).

See attached Supplementary Information for further figures, movies, and methods.

## For Table of Contents Only

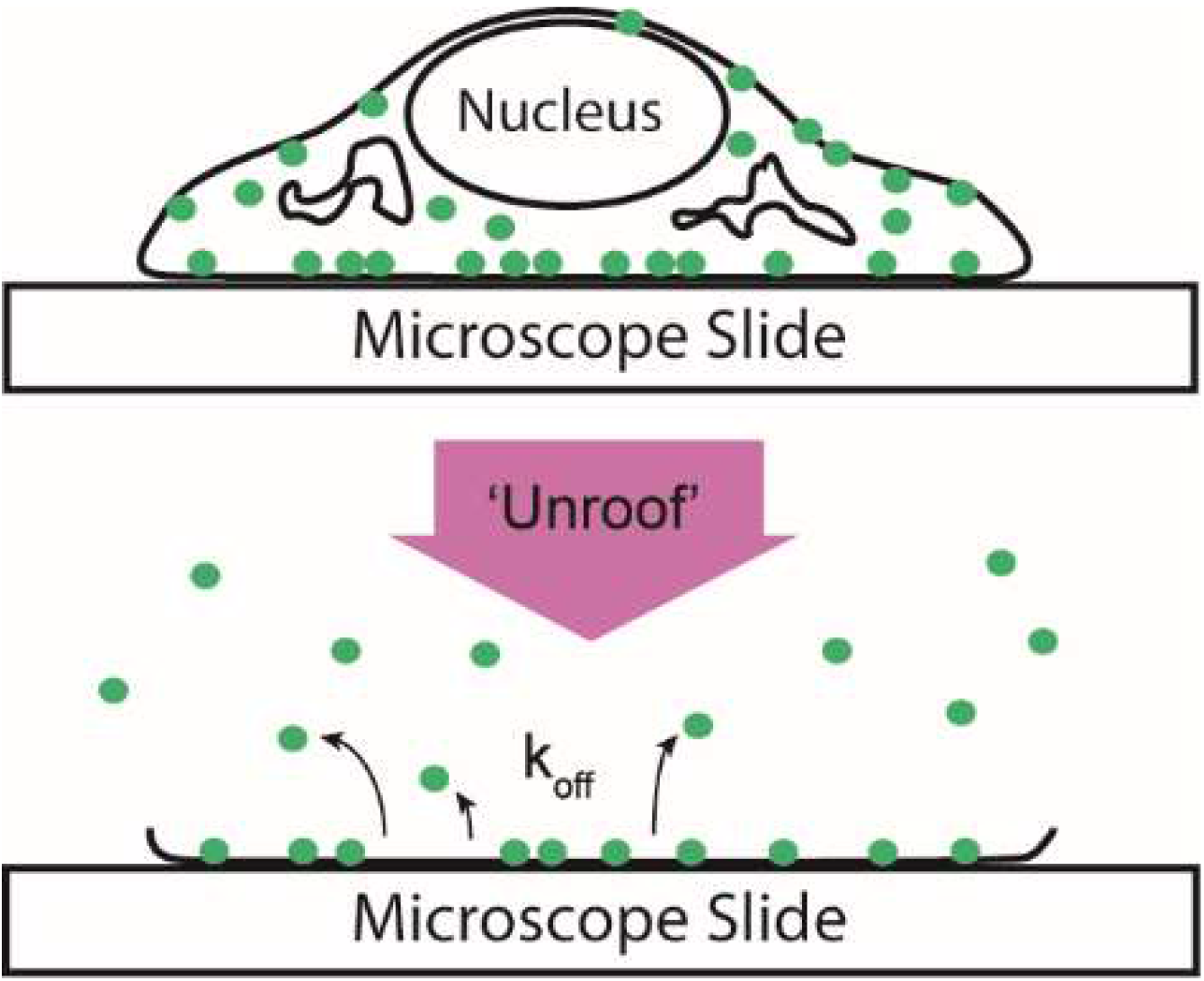

