## Supplementary Information for "Phosphatidylinositol phosphate binding domains exhibit complex dissociation properties at the inner leaflet of plasma membrane sheets"

Supplementary Figures

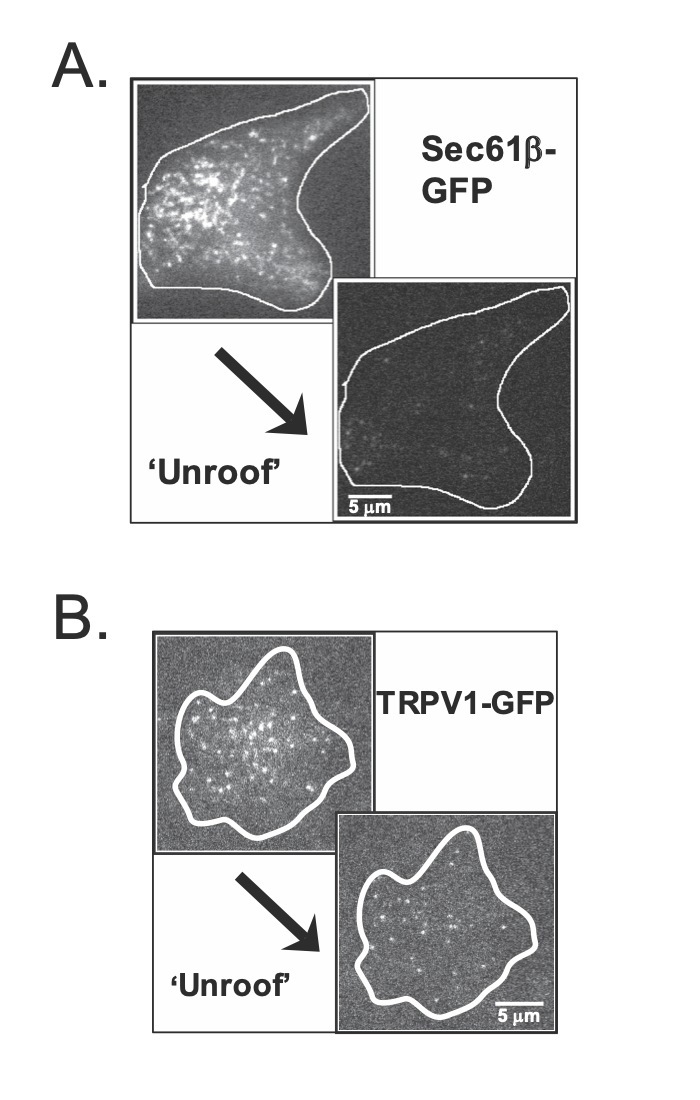

Supplemental Figure 1. Cell unroofing isolates plasma membrane (PM)-bound proteins with low endoplasmic reticulum (ER) contamination. (A) Imaging with TIRF microscopy shows that after unroofing, HEK293T cells expressing the ER marker Sec61B-GFP exhibit a significant loss of intracellular membrane across the cell footprint compared to the intact cell. (B) A HEK293T cell (outline in white) expressing the integral membrane protein TRPV1-GFP has diffuse background fluorescence. Individual TRPV1-GFP channels residing in the PM remain after unroofing while the contaminating signal from ER pools of TRPV1-GFP is no longer present.

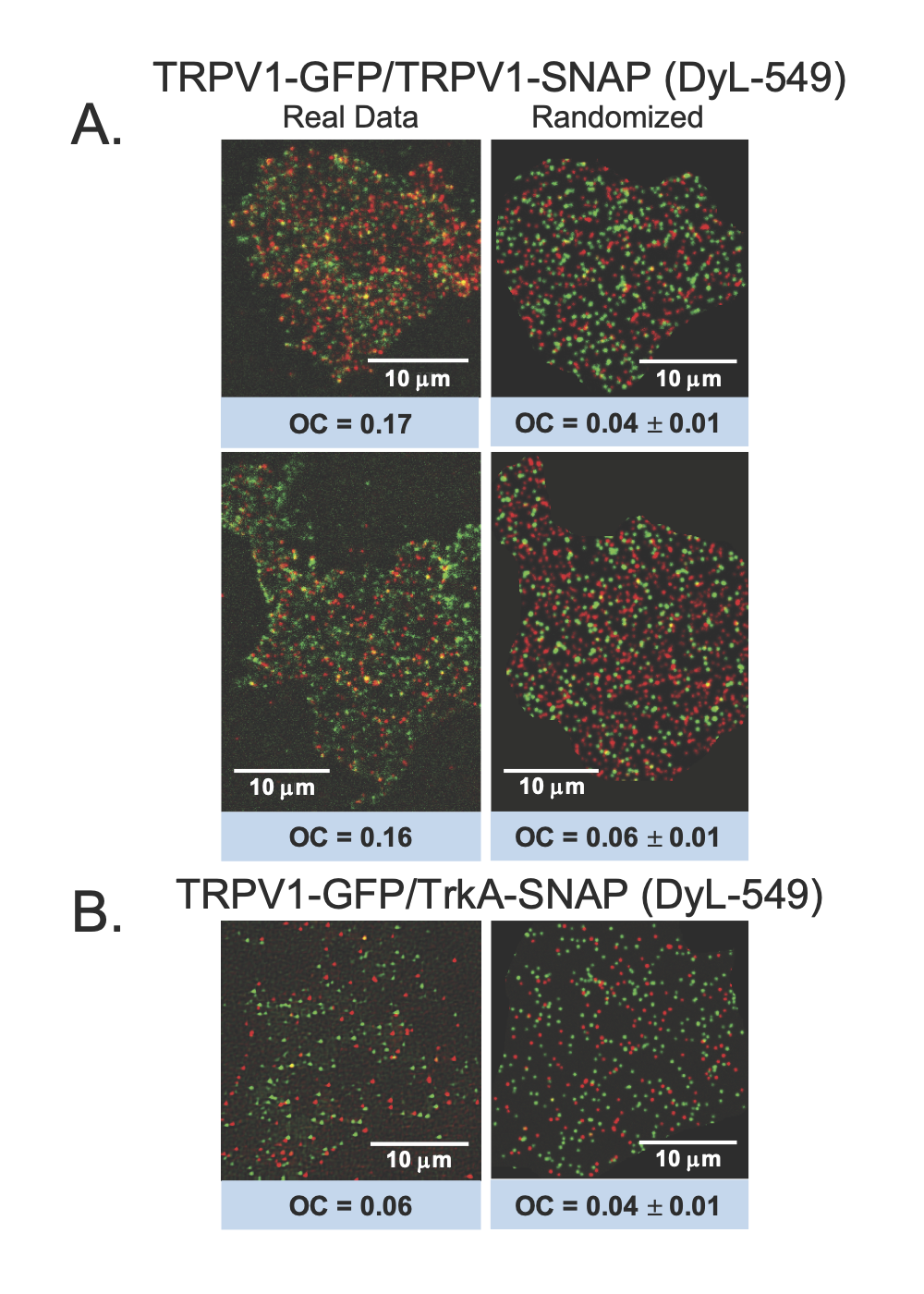

Supplemental Figure 2. TRPV1-GFP and TRPV1-SNAP colocalize. (A) Two unroofed HEK293T cells expressing TRPV1-GFP and TRPV1-SNAP labeled with DyL-549 (left). In comparison to mean object colocalization (OC) values derived from randomized simulation of the two fluorophores (right, n=5), the unroofed cells consistently show a higher coincidence of the two fluorescently labeled TRPV1 constructs (see methods). (B) An unroofed cell expressing TRPV1-GFP and TrkA-SNAP labeled with DyL-549. Overlap of the PM-bound fluorescent proteins evaluated by OC (left) is not easily distinguishable from randomized simulations of the fluorophores (right, n=5).

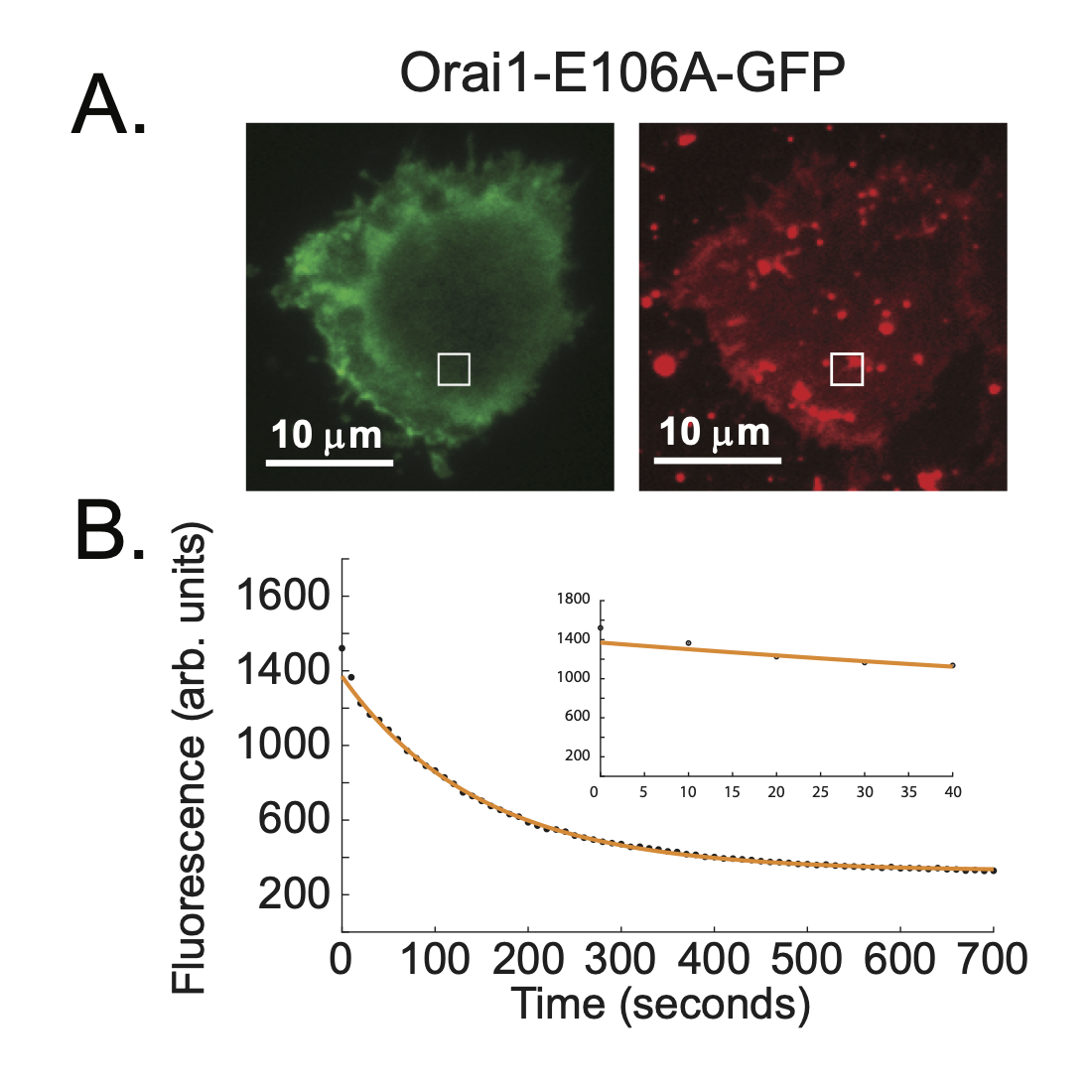

Supplemental Figure 3. Photobleaching occurs at a significantly slower rate than that measured for dissociation of pleckstrin homology (PH) domains from the membrane. (A) HEK293T cells expressing Orai1-E106A-GFP imaged with TIRF microscopy were excited at 488 nm to follow photobleaching (left) and excited at 561 nm with DiI to confirm the presence of PM sheets (right). White boxes demarcate the selected region of interest (ROI). (B) Representative fluorescence trace specific to the ROI. Cells were unroofed 10 s into each recording, with fluorescence decline corresponding to photobleaching of the ion channel over the course of 700 s. Inset displays the fluorescence in the first 40 s of imaging (corresponding to the length of time for which cells expressing PH domains were imaged). Orai1-E106-GFP k_PB_ = 0.0072 ± 0.0009 s^-1^.

Supplementary Movies

Supplemental Movie 1: Loss of fluorescence in AKT-PH-GFP expressing cell following unroofing. Single frame and fluorescence data is shown in Figure 1A,B.

Supplemental Movie 2. Loss of fluorescence in PLC-δ1-PH-GFP expressing cell following unroofing. Single frame and fluorescence data is shown in Figure 1A,B.

Supplemental Movie 3. Rebinding of PLC-δ1-PH-GFP onto plasma membrane sheet of neighboring cell following unroofing. Imaging sequence corresponds to Figure 4.

Supplemental Movie 4. Single molecule fluorescence imaging of PLC-δ1-PH-GFP interacting with plasma membrane sheet of unroofed cell. IP_3_ added to the perfusion chamber at 6 second mark. Image sequence corresponds to Figure 5.

Supplemental Movie 5. Loss of fluorescence in PLC-δ1-PH-GFP expressing cell following unroofing while in the presence of 100μm IP_3_. Single frame and fluorescence data is shown in Figure 6A,B.

Supplementary Methods

DNA and Molecular Biology

The GFP derivative EGFP was used in all fluorescently-tagged constructs and is simply referred to as GFP throughout the text ^1^. Constructs of PLC-δ1-PH-GFP and AKT-PH-GFP are fusion proteins with the GFP located on the N-terminus of the PH domain. Expression of the PLC-δ1-PH-GFP constructs was from the pUC plasmid under the control of the CMV promotor. Expression of the AKT-PH-GFP constructs was from the pcDNA3.1 plasmid under the control of the SV40 promoter. Site-directed mutagenesis was performed on the PLC-δ1--PH-GFP and AKT-PH-GFP constructs to introduce the A207K mutation, which produces a dimerization deficient form of GFP ^2^. TRPV1-GFP, TRPV1-SNAP, and TrkA-SNAP are fusion proteins of wild-type rat TRPV1 or wild-type rat TrkA with the GFP or SNAP domain located on the C-terminus. Expression of the TRPV1 and TrkA constructs was from the pcDNA3.1 plasmid under the control of the CMV promoter. All constructs were acquired from the lab of Sharona Gordon (Univ. of Washington, Seattle WA).

Cell Culture and Transfection

HEK293T/17 cells (CRL-11268 ATCC, Manassas, VA) were cultured at 37°C and 5% CO_2_ in Dulbecco’s modified Eagle medium (DMEM, Gibco ThermoFisher, Waltham MA) containing 25 mM D-glucose, 1 mM sodium pyruvate, and 4 mM L-glutamine. Culture medium was supplemented with 10% fetal bovine serum (FBS, Gibco, ThermoFisher, Waltham MA) and penicillin/streptomycin (Gibco, ThermoFisher, Waltham MA). Cells were plated on 100 mm dishes at a density chosen to achieve 30–50% confluency at the time of transfection. Cells were transfected using Lipofectamine 2000 (Gibco, ThermoFisher, Waltham MA) according to the manufacturer’s instructions with the following modifications: for each plate, a total of 3 μg DNA and 6 μl Lipofectamine in 600 μl Opti-MEM (Gibco, ThermoFisher, Waltham MA) was used. After incubating for 16–22 hours, cells were passaged onto coverslips for unroofing and the medium was replaced with HEPES buffered Ringers (HBR, 140 mM NaCl [Sigma-Aldrich, St. Louis MO], 10 mM HEPES [Sigma-Aldrich, St. Louis MO], 5mM D-(+)-glucose [Sigma-Aldrich, St. Louis MO], 4 mM KCl [Sigma-Aldrich, St. Louis MO], 1.8 mM CaCl_2_-2H_2_O [Sigma-Aldrich, St. Louis MO], and 1 mM MgCl_2_-6H_2_O [ThermoFisher, Waltham MA]; initial pH measured at 5.2 and adjusted to 7.4 with NaOH [ThermoFisher, Waltham MA]; filter sterilized through a 0.45-μm membrane). Coverslips for unroofing experiments were prepared by incubation with filter sterilized 0.1 mg/ml poly-D-lysine hydrobromide (P7886, MW 30,000-70,000, Sigma-Aldrich, St. Louis MO) in phosphate buffered saline (PBS, pH 7.4, Sigma-Aldrich, St. Louis MO) for 10 min and were then washed using PBS to remove excess poly-D-lysine.

Single Molecule Fluorescence Imaging

For single molecule imaging, TIRF microscopy settings were altered to visualize individual fluorophores at the membrane; the EM gain was increased from 50 to 300 and the laser excitation light was increased. Single molecule imaging was conducted by unroofing as described above except that the perfusion system was stopped immediately after unroofing to retain free PLC-δ1-PH-GFP in solution. After removing the probe from the chamber, we imaged the plasma membrane sheet to record binding and unbinding events at the single molecule level. We also introduced IP_3_ into the chamber to inhibit the binding of PLC-δ1- PH-GFP to PIP_2_ in the membrane sheet as follows: 6 s after imaging commenced, 4 μL of 10 mM IP_3_ was added to the bath solution in the chamber for a final concentration of 100 μM IP_3_; this was repeated every 6 seconds for a total of 4 times, reaching a final concentration of 400 μM IP_3_ (Supplemental Movie 5). Unroofing was confirmed as described above.

AFM imaging:

Freshly unroofed cells were treated with 0.05% glutaraldehyde for 10 minutes followed by repeated (3x) rinsing and storage in SB at 4°C. Within 3 hours, samples were transferred to an AFM imaging facility, where images were acquired by PeakForce tapping mode with a Bruker AFM system (Bruker Corporation, MA USA). Images were processed for rendering by Gwyddion (www.gwyddion.net)

Image processing and fluorescence data fitting:

Image analysis was performed with ImageJ ^3^. We implemented the JACoP plug-in (https://imagej.nih.gov/ij/plugins/track/jacop.html) to acquire optical colocalization (OC) measurements for Figure S2, and randomized puncta of fluorescence were generated with the model feature of the GDSC plug-in (<https://sites.imagej.net/GDSC-SMLM/>) ^4^. For fluorescence unroofing measurements, regions of interest were created by selecting two 10x10 pixel representative samples for each individual cell. Background fluorescence was corrected for by defining baseline (zero fluorescence) as the minimum fluorescence value in a given dataset. Data was optimized for bulk fitting by truncating fluorescence data to remove initial fluorescence drop-off following unroofing attributable to the rapid shedding of cytoplasmic PH domains from the unroofed cell. Fluorescence data corresponding to the 10 s following this initial drop-off were analyzed and plotted with Matlab (Mathworks, MA USA) and fit with the following single and double exponential functions:

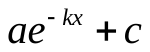

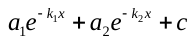

Graphics comparing our compiled unroofing measurements were generated using the Data Analysis with Bootstrap Estimation (DABEST) package in Matlab ^5^.
